# Contingency, Repeatability and Predictability in the Evolution of a Prokaryotic Pangenome

**DOI:** 10.1101/2023.03.20.533463

**Authors:** Alan Beavan, Maria Rosa Domingo-Sananes, James O. McInerney

## Abstract

Pangenomes exhibit remarkable variability in many prokaryotic species. This variation is maintained through the processes of horizontal gene transfer and gene loss. Repeated acquisitions of near-identical homologs can easily be observed across pangenomes, leading to the question of whether these parallel events potentiate similar evolutionary trajectories, or whether the remarkably different genetic background of the recipients mean that post-acquisition evolutionary trajectories end up being quite different. In this study, we present a machine learning method that predicts the presence or absence of genes in the *Escherichia coli* pangenome based on the presence of other accessory genes within the genome. We are, in effect, asking whether gene acquisitions potentiate similar evolutionary trajectories or not. Our analysis leverages the repeated transfer of genes through the *E. coli* pangenome to observe patterns of repeated evolution following similar events. The presence or absence of a substantial set of genes is highly predictable, from other genes alone, indicating that selection potentiates and maintains gene-gene co-occurrence and avoidance relationships deterministically over long-term bacterial evolution despite differences in host evolutionary history. We propose that the pangenome can be understood as a set of genes with relationships that govern their likely cohabitants, analogous to an ecosystem’s set of interacting organisms. Our findings highlight intra-genomic gene fitness effects as key drivers of prokaryotic evolution, with ensuing pangenome-wide emergence of repeated patterns of community structure.

## Introduction

Evolution by horizontal gene transfer (HGT) and differential loss causes remarkable variation in gene content in bacterial genomes, both within and between populations (Tettelin et al., 2005, Treangen and Rocha, 2011, McInerney et al., 2017, Puigbò et al., 2014, Vos et al., 2015). Genes present in all genomes in a population constitute the core genome, while genes that are found only in some lineages are accessory genes. The union of these two sets make up the pangenome. Intra-specific HGT, mediated by plasmids, phage, and transduction account for most gene transfers into a genome. Though there has been some disagreement on the relative influences of random drift and natural selection on structuring pangenomes, a body of work has now shown that the presence or absence of specific genes (genetic background) can influence the presence or absence of others (Rosconi et al., 2022, Beavan and McInerney, 2022, Caudal et al., 2022). Consequently, the content of every contemporary prokaryotic genome is an outcome of its history of vertical and horizontal gene transmission and has emerged *via* a combination of internal (intragenomic) and external (ecological) fitness effects (McInerney, 2023) in addition to stochastic, non-adaptive evolution (genetic drift).

It is also unclear how evolutionary responses, say to the acquisition of a gene by horizontal gene transfer, are sensitive, or robust, to differences in evolutionary history. In his book, “Wonderful Life: The Burgess Shale and the History of Nature” (1990), Stephen J. Gould set out a thought experiment where the “tape of evolution” could be replayed from any point in history. He proposed that because evolutionary trajectories were contingent on unpredictable events then no two runs of the tape would lead to the same outcome. Many recent studies have suggested this view is too rigid. Experiments designed to mimic replaying of the tape, such as parallel evolution experiments, have suggested that historical contingency does indeed have an effect but that some aspects of evolution are deterministic – *i.e*. they are likely to happen each time we replay the tape (Blount et al., 2018, Lenski, 2017, Losos, 2010, Leiby and Marx, 2014, Wolfe et al., 2021). Until now, it has not been obvious how, or even if, the contingency-deterministic question relates to prokaryote genome evolution. In prokaryote pangenome evolution, repeated HGT can introduce homologs of the same gene family into divergent genomes that contain unique but overlapping sets of genes. The incorporation of these genes into different genetic backgrounds, allows us to address the contingency-determinism question through retrospective analysis of the subsequent outcomes. We identify a deterministic outcome if all, or most, recipient lineages evolve in similar ways after gene acquisition, while the alternative is that prior events – divergence in gene content of the recipient genomes – would play the more important role, and post-acquisition evolution of the different lineages would therefore be markedly different.

In detail, we define a deterministic evolutionary trajectory as the acquisition of a gene that in turn potentiates the acquisition, avoidance, retention, or loss of one or more other genes. In other words, certain evolutionary outcomes are highly likely, thanks to the influence of intragenomic selection on genotypes. Repeated acquisition and loss of a gene, while necessary, is insufficient to imply deterministic evolution. Hallmarks of determinism include the emergence of repeated biases in gene content, including the selective recruitment of another gene, or selective loss of another gene, following horizontal transfer. Owing to the prominent role of stochastic processes in evolution, it is unlikely that gene content evolution is entirely deterministic or entirely driven by contingency, but rather it falls somewhere on the spectrum between both extremes. The question is which end of the spectrum is closest. Today, several thousand complete prokaryotic genomes are available, providing enough data to address the issue. Therefore, we can ask whether a gene’s presence or absence in a genome is predictable, based solely on the gene content of the rest of the genome, implying deterministic evolution, or unpredictable, implying contingent events significantly influence evolutionary outcomes.

Several programs have been developed to find co-evolving gene pairs and to infer co-evolving modules (Lassalle et al., 2019, Cohen et al., 2013, Whelan et al., 2020, Harling-Lee et al., 2022, Hall et al., 2021, Whelan et al., 2021). However, if gene presence or absence in a genome is indeed affected by other genes, then there is likely to be a complex combination of positive and negative intragenomic effects that extend beyond pairwise correlations. To incorporate these more complex and subtle patterns, we used a Random Forest approach (Ho, 1995). Random Forests aggregate information from individual decision trees, which themselves summarise the conjunction of features that lead to predictions of gene presence or absence. In our case, the features of the Random Forest analysis were the genes in the pangenome, encoded in presence-absence vectors.

Random Forests employ combinations of genes, not just pairwise comparisons to test gene predictability in quite different genetic backgrounds. Therefore, complex conditional relationships between sets of genes can be revealed when the random forest approach is used. Furthermore, the random forest approach can assess whether inferences are generalisable. The model that we use to predict gene presence or absence is parameterised on a training dataset and evaluated on a test dataset (Ho, 1995). If the model built using the training dataset does not describe the patterns found in the test set, it is probable that the pattern is an artefact of the training set, and the model should be considered inadequate. However, if the model makes accurate predictions in the test set, it appears to describe general properties of the entire dataset. Finally, Random Forest models make predictions in a directed manner, where one gene might predict the presence or absence of another, meaning we can say whether a gene is predictable and if so, we can also identify its predictors.

In this paper we demonstrate that a substantial proportion of *Escherichia coli* accessory genes can be predicted by the other genes in the genome. *E. coli* has a large accessory genome (Welch et al., 2002, Lukjancenko et al., 2010), and occupies a wide range of niches (Kaper et al., 2004). The *E. coli* pangenome has evolved divergent gene content over time – so much so, that a gene that is horizontally transferred from one *E. coli* to another will often find itself in a considerably different ensemble genetic background. We have analysed the predictability of gene content evolution following the repeated transfer of genes into these diverse genetic backgrounds. This is a natural equivalent of what Blount et al. (2018) called a “historical difference experiment”. We have typified the effects of accessory genes’ presence on the presence or absence of other genes into three categories typically used by macroecologists to describe interactions between species *sensu* McInerney, 2023). We define *Mutualism* as both genes predicting each other’s presence where each gene has a similar effect on the probability of finding the other. *Commensalism* refers to the situation where one gene strongly depends on the presence of another, but the reverse dependence is much weaker or non-existent. *Competition* is when two genes appear to avoid being in the same genome. Note that we are not ascribing behaviours to genes, these categories are simply descriptions of patterns.

## Materials and Methods

*E. coli* genomes were downloaded from the NCBI genome database (Sayers et al., 2022) using the NCBI command line utility “datasets” 12.17.2 accessed 01/05/2022. All annotated *E. coli* genomes were downloaded provided they were of the highest completeness level (complete). If a genome had been assembled by both the GenBank and RefSeq methods, the GenBank (GCA) annotation was maintained, however, if only a RefSeq (GCF) annotation was in the database, this was retained. The final dataset consisted of 2,341 genomes. The full list of accession numbers is included in supplementary list 1.

All genomes were re-annotated using PROKKA version 1.14.6 (Seemann, 2014), using the annotation mode “bacteria”, an e-value cutoff of 10^-9^ and a minimum query coverage of 80% were required when assigning functional annotations. The *E. coli* pangenome was inferred using Panaroo version 1.2.9 (Tonkin-Hill et al., 2020) with the mode set to sensitive. It was important to include rare genes, owing to the possibility that they would have important effects on the prediction of other genes.

The gene presence-absence matrix produced by Panaroo was processed so that genes present in more than 99%, or less than 1% of genomes were removed. The matrix was further modified by converting gene names to “1” and empty fields to “0” so that 1 represented presence and 0 represented absence. In addition, genomes with identical patterns of gene presence and absence were collapsed into the same vector. Genes with identical presence-absence patterns (PAPs) across our genome sample were also collapsed into gene family groups (Supplementary table 1). Essentially, this means that both the features and predicted variable in these analyses are PAPs rather than genes *per se*, most of which are represented by only one gene but some of which are represented by multiple genes. The list of genomes with non-unique gene repertoires is found in supplementary table 2.

To minimise the impact of phylogeny on understanding predictability, repeatability, and contingency, we impose the requirement that we only study genes whose distribution is not “clumped” on one or a few clades. A backbone phylogeny was inferred in order to evaluate the distribution of gene content across the tree. Alignments of universal single-copy genes were constructed using MAFFT version 7.490 (Katoh et al., 2002). The resulting 337 gene alignments were concatenated to form a superalignment and a maximum likelihood phylogeny was reconstructed using IQTree 2.2.0 (Nguyen et al., 2015) with extended model selection (Kalyaanamoorthy et al., 2017) and 1,000 ultrafast bootstrap replicates (Hoang et al., 2018). The tree was rooted at the midpoint. To assess the distribution of each gene on this tree we calculated Fitz and Purvis’ D statistic (Fritz and Purvis, 2010), retaining only those pairs of genes where the target nodes have a D statistic greater than zero, though source nodes could still have D < 0. To contextualise the meaning of the D statistic we also calculated the parsimony score for each gene using PAUP v4.0a (Swofford, 2002). This metric represents the minimum number of times a gene has shifted from present to absent or absent to present along the phylogeny. Filtration of the edges was carried out using Sqlite3 (Hipp, 2020). In practice, we only study genes that have changed character-state from present to absent or *vice versa* on the tree at least 8 times, and using the D-statistic (Fritz and Purvis, 2010), we insist on only analysing genes where D is > 0.

A model of gene presence or absence was inferred using random forests. Prediction models for each PAP are calculated separately, meaning the number of times that our random forests were trained was equal to the number of PAPs in the processed matrix. For each PAP being predicted, the genomes are randomly split into a training set, equal to 75% of the genomes and a test set of the remaining 25%. These sets were stratified by the gene presence absence state being predicted, meaning that the proportions of genomes with any given gene being present or absent, remained approximately the same in the training and the test datasets. These test and training datasets were assigned independently during the prediction of every PAP in every analysis. Decision trees were then generated by taking a random sample of genes with size equal to the square root of the number of genes in the dataset and the gene that most evenly split the training set out of this subset formed the first node in the tree (according to Pedregosa et al., 2011). This process was repeated on each side of each decision node until either the maximum depth was reached, or the remaining sample of genomes all had the same state for the gene in question. The effect of variation in the number of trees generated, and the maximum depth of those trees was empirically evaluated according to the average performance of the analysis in predicting the presence or absence of each gene (Supplementary Figure 1). We chose the maximum depth and number of trees above which the performance of the models on the test set did not increase substantially. For the analyses in this manuscript, including those where the dataset was downsampled, we used 1,000 trees with a maximum depth of 16 nodes per tree.

For each gene (or set of genes with the same PAP), a prediction of either its presence or its absence is obtained for each genome in the test set according to the model generated using the training set. For each gene, 4 performance metrics were taken. These were Accuracy, Precision, Recall and F1 score, all defined according to (Van Rijsbergen, 1979). As our classification algorithm has two classes so we record a version of recall, precision and F1 score separately for both the gene presence and for the gene absence classes.

In addition to performance statistics, the Gini importance (Breiman et al., 1983) of each gene in predicting each other gene was added to an *n* by *n* matrix where *n* is the number of PAPs in the dataset. Here, the Gini importance is the contribution of a predictor PAP in separating the dataset into genomes where the test gene is present and those where it is absent averaged over all trees in the random forest. This Gini importance value can be used as a measure of the strength of the impact of the predictor gene on the presence/absence state being predicted. Use of this statistic in this way is appropriate where the predictor variable has comparable numbers of classes (Nembrini et al., 2018). All machine learning algorithms were implemented using the scikit-learn python module version 1.0.1 (Pedregosa et al., 2011). Code is available at: https://github.com/alanbeavan/pangenome_rf (Supplementary Figure 2).

To generate a null expectation of performance and importance, and to evaluate how often we expect to see associations arise by chance, we randomly permuted the gene presence-absence matrix, keeping the number of genomes in which each gene is found. Results obtained using the original pangenome were compared to the permuted datasets. Finally, to test the repeatability of the results, analyses were performed 100 times.

### Post-processing the network

Initially, all relationships between PAPs with a GINI importance of less than 0.01 were removed from each network examined. This threshold is arbitrary, chosen to restrict the number of gene-gene relationships to the most highly ranked, a number that could be visualised. All importance values above the cutoff threshold were significantly greater than those generated by randomly permuted data (Supplementary Figure 3). After examination of the networks resulting from several different thresholds, ours was chosen because it generated networks that could be easily visualised yet described enough predictive relationships to provide an understanding of pangenome evolution.

Co-occurrence is distinguished from avoidance by comparing the frequency of the in genomes where the source is present with the frequency of the target full dataset. If the frequency of the target is higher when the source is present, the relationship is set as “co-occurrence”. If it is lower, then the relationship is flagged as “avoidance”. To ensure our model only contained strongly predicted relationships we kept only those edges where the target node was classified as accurate, which here means the F1 score for both classes (present and absent) was greater than or equal to 90% in the test set.

Artefactual splitting of *de facto* gene families can, in principle, take place during pangenome construction. Therefore, using BLASTn (Altschul et al., 1990, Altschul et al., 1997), we subjected the sequences of each gene family involved in any apparent avoidance relationship to a comparison with the gene family they avoid. A total of 563 gene families (before filtering for performance and D) featured at least one sequence that was identical to a sequence placed in a different family. Gene families with this property were therefore removed from our results. After this step, any two sequences from avoidant gene families never share identity of at least 50% of their nucleotides or produce a significant BLAST hit with an associated E-value < 10.

### Functional classification of genes and enrichment analysis

Eggnog mapper version 2.1.8 (Huerta-Cepas et al., 2017) was used to assign gene ontology terms (go-basic release 2022-07-01, Gene Ontology Consortium, 2021, Ashburner et al., 2000) and KEGG pathway (Kanehisa et al., 2016, Kanehisa and Goto, 2000) terms for each gene that included in this analysis (those with between 1 and 99% genome occupancy). Diamond-BLAST (Buchfink et al., 2021) was used to search the eggnog mapper database for sequences with an e value <10^-5^. For functional enrichment analysis, the program find_enrichment.py which is part of the GOa tools suite (Klopfenstein et al., 2018).

### Calculating physical linkage between genes

The physical distance between two genes was measured in the number of genes they are separated by, rather than base pairs. The position of each gene of the pair was extracted in each genome from the Panaroo output (Tonkin-Hill et al., 2020). For each pair of genes, the distance between was defined as the minimum of distance between the 2 genes going clockwise and going anti-clockwise around the circular chromosome.

### Categorising accessory gene dynamics

We placed observations of gene associations into three mutually exclusive categories. Firstly, mutualisms, where the presence of one gene predicts the presence of another. For mutualistic relationships where the presence of one gene predicts the presence of the other, we require that the strength of the prediction is similar in both directions, with the F1 statistic for both classes greater than or equal to 0.9 and the D statistic is greater than zero. For commensalisms where the presence of one gene is highly dependent on the presence of another, but the reciprocal dependency is either far weaker or non-existent, the more abundant, or “host gene”, must be present in almost all (>= 99%) genomes where the putative “commensal” is present. Additionally, the proportion of genomes without the commensal where the host is present must be at least 20% of the proportion of genomes in the full dataset containing the host. That is, if the host gene occupies 50% of the genomes in the full dataset, it must be present in 10% of the genomes where the commensal gene is absent. Otherwise, we do not classify a relationship as commensal. If a gene-gene association was classified as commensal it could not be classified as mutualistic. Finally, we consider two genes to be in competition or antagonistic of one another when the absence of one gene is strongly predicted by the presence of the other or *vice versa*.

### Data visualisation

Gephi v0.10 (Bastian et al., 2009) was used to visualise networks. Network layout was achieved using the Fruchterman-Reingold layout algorithm (Fruchterman and Reingold, 1991). Other graphics were produced using custom python or R scripts.

## Results

### A substantial subset of accessory genes in *E. coli* can be predicted accurately

The *E. coli* pangenome inferred in this study contained accessory gene families with 12,841 unique PAPs that were present in more than 1% and less than 99% of genomes and were hence included in this study. The presence or absence of 3,922 (30.5%) PAPs could be accurately predicted (both F scores >= 0.9) in the test set after the random forest model had been trained. From this accurately predicted dataset, a total of 2,144 (54.7%) had an associated D-statistic greater than 0, meaning they were distributed widely on the tree. The remaining 1,778 PAPs were “clumped” on the tree and therefore it is more difficult to ascribe causality to their association, when a very good explanation might be that they were simply acquired at more-or-less the same time and have been vertically inherited together since then. Figure S1 shows that although the D score is not directly proportional to parsimony score, it correlates strongly, meaning that all 2,144 PAPs with a D score of greater than or equal to zero also had a parsimony score of at least eight though most had a much higher score (Supplementary Figure 4). This means that we have only examined relationships in the data where genes have been acquired and/or lost at least 8 times across the pangenome, and furthermore, we insist that their distribution is widespread and not localised (Fritz and Purvis, 2010). We focus on this set of 2,144 PAPs because they manifest a broad, patchy distribution across the phylogeny, stemming from a combination of lateral gene transfer and loss, and we can accurately predict their presence or absence based on the other genes present in the genome. Node ranking was assigned according to the PageRank (Brin and Page, 1998) algorithm weighted by Gini importance of each incoming arc. A considerable amount of variation was observed in the weight ascribed to the nodes on this network when we employed the PageRank approach to ascertain node importance. Specifically, to attach a ranking to a node, the algorithm combines the number of incoming arcs, the weight of the arcs and the importance of the source nodes (Brin and Page, 1998). Node size in figure 1 is proportional to node rank, as judged by the PageRank algorithm.

**Figure 1:**
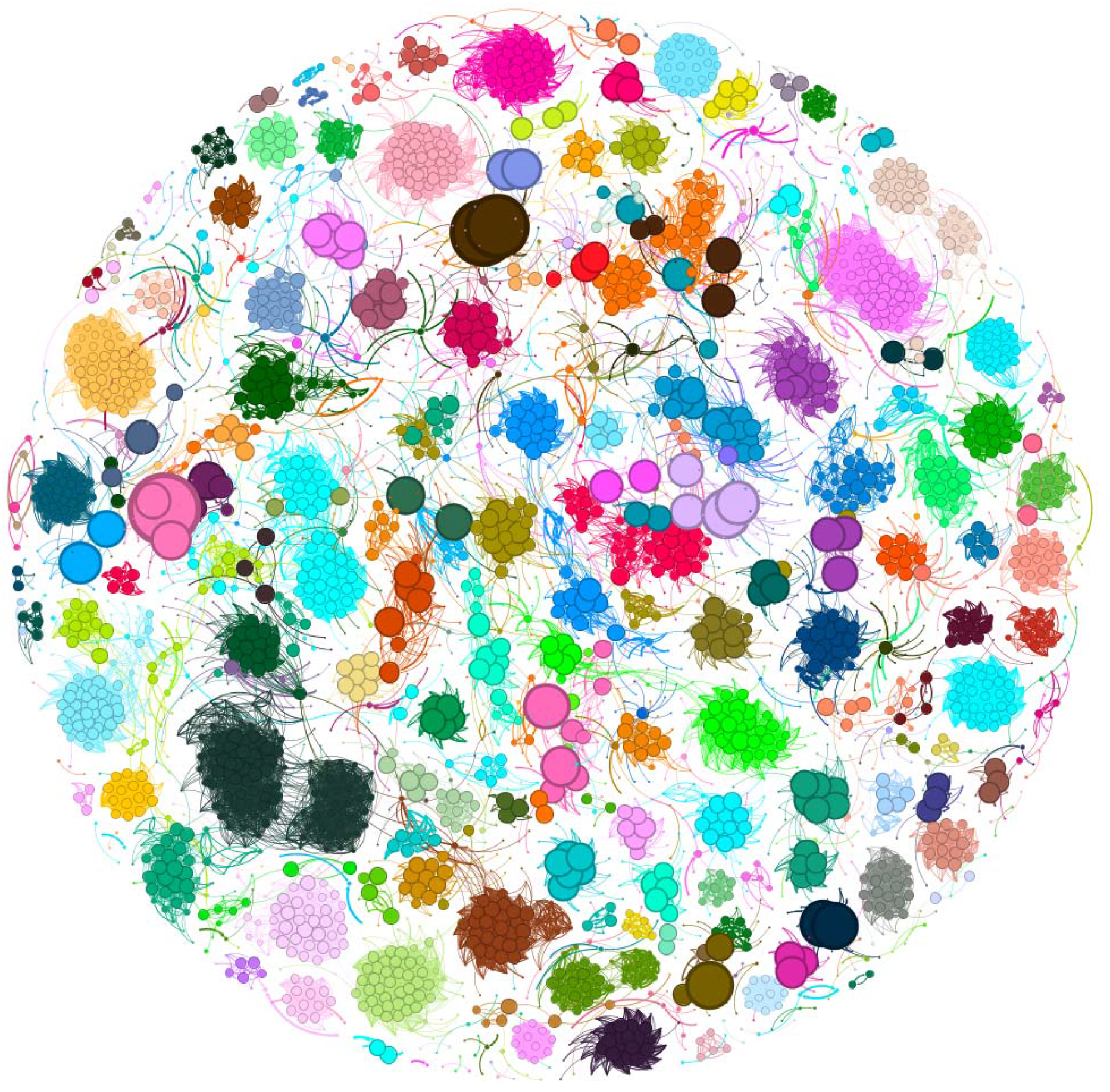
The coincident relationships of predictable genes and their predictors. The nodes are gene families, or groups of gene families with the same PAP, and the edges are coincidence relationships with the arrow pointing at the node whose presence is predicted by the other. Edge thickness is proportional to the importance value, while node size is proportional to the PageRank (Brin and Page, 1998) value for that node. Node colour indicates community as identified by the Louvain algorithm (Blondel et al., 2008).

To evaluate whether the presence absence matrix of 12,841 uniquely distributed gene families is no better structured than random expectation and that ostensible gene predictability simply arises by chance in datasets of this nature, we compared results from the original data to those recovered from a permuted version of the dataset with the same size and composition. We shuffled the presence absence pattern of each gene family to induce breakage of any correlation across patterns, while preserving individual gene frequency and the distribution of frequencies across the dataset. The resulting randomised dataset was used as input to the same model generation process as the unpermuted data. Not one of the models from this randomised dataset generated predictions for even a single gene with an F1 statistic for both classes above our threshold. In addition, we observed a total of zero Gini importance values greater than 0.01 for the randomly permuted dataset. The unperturbed data, in contrast, resulted in >2000 accurately predicted PAPs each time it was run. Accordingly, we can reject the hypothesis that these empirical observations of associations have arisen due to chance in our dataset, or that the structure of the pangenome dataset has no more gene-gene correlations than the structure of randomly assembled data.

To investigate the effect of data quantity on gene predictability, we carried out a sensitivity analysis on dataset size. We randomly eliminated 50%, 75%, 90% and 95% of the genomes in the dataset then repeated our random forest prediction 10 times per dataset. In each case, reducing the number of genomes substantially and significantly reduced the number of PAPs that were accurately predicted, while having a much smaller effect on the number of total PAPs that could be analysed. For example, the average number of accurately predicted PAPs, over 10 repeated analyses after filtering PAPs with D score <0, using 50% of the genomes was 1,650/12,642 (13.1%) compared with the 2,144/12,841 (16.7%) in our full analysis. When only 5% of genomes were included in the study, an average of 713/11,644 (6.1%) PAPs were predicted accurately (Supplementary Figure 5).

The links between the 2,144 predictable PAPs, were used to construct a network with 33,426 edges featuring all well-predicted target nodes and their predictors (Figure 1). This network consisted of 243 connected components ranging in size from 2 to 248 nodes, featuring both coincident and avoidance edges *sensu* Whelan et al. (2020). By considering only the coincident relationships (33,138 out of 33,426 edges) we found 240 connected components containing between 2 and 244 nodes. Taking only avoidance relationships, 28 connected components were generated with a range from 2 to 22 nodes in size. As non-unique gene patterns are collapsed into one entity, both in the analysis and presentation of results, some of the nodes represent multiple genes. 827 of the well predicted PAPs applied to multiple genes. In total, independent of whether they are well predicted by our random forest model or not, 19,137 genes had non-unique PAPs and were collapsed into 4,172 patterns that were then used both as features for prediction and as patterns to predict.

Owing to the stochastic nature of the Random Forest approach, we repeated the analysis 100 times, each time splitting the data into training and test sets differently. Out of the 12,841 accessory genes with unique PAPs analysed, 3,137 were never classified as predictable, 2,222 were always classified as predictable (before filtering by D score) and the remaining 4,831 were classified as predictable in only some analyses (Supplementary Figure 6). 870 of the PAPs that were always predictable had a D score >0. To understand what makes a gene’s presence or absence predictable, we compared these consistently predictable PAPs to the least predictable gene families in the dataset, which we defined as the set of genes that never passed our thresholds of predictability in any of the 100 iterations of the RF algorithm. To ensure a balanced comparison, we took the 870 PAPs with the lowest average combined F1 score over the 100 repeats to generate two equally sized datasets representing the most, and least predictable genes.

After functional annotation using EGGNOG mapper (Huerta-Cepas et al., 2017) gene ontology (Gene Ontology Consortium, 2021) and KEGG (Kanehisa et al., 2016) pathway enrichment analyses were performed. Using p-values that were corrected by false discovery rate and other conventional methods including Bonferroni, the set of most predictable genes were not enriched in any function, cellular location, or process, and were not enriched in any gene ontology terms in either biological process, molecular function, or cellular compartment. Additionally, only two KEGG pathway terms were enriched in the consistently predictable genes. These were “bacterial secretion system” and “flagellar assembly”.

Amongst the 870 least predictable genes, 3 Biological Process terms were enriched and 26 were underrepresented, while for the cellular compartment terms, 3 were enriched and 7 were underrepresented, and for molecular function, there were no enriched terms, while 3 were underrepresented. No KEGG pathway was found to be enriched in this data set, after FDR correction. In total, 6 GO terms were enriched in the low predictability set of genes, while 36 GO terms were significantly underrepresented (Supplementary Table 3).

Given that the most predictable genes appear to contribute to a range of functions, we investigated the extent to which physical linkage determined their predictability. We compared the positions of the genes that shared an association and measured the distance between them (in numbers of genes rather than base pairs) as inferred by Panaroo. From these distances, clearly linkage plays an important role in the association of two genes (Supplementary Figure 7). 68.7% of pairs of associated genes in the same genome are separated by 10 genes or fewer, in the set of most well predicted genes. However, even in the most predictable set of genes, there are several occurrences of genes that are not closely physically linked with the genes that they are coincident with (10.4% of pairs of genes were separated by at least 21 genes), so it cannot be the only factor at play. In addition to pairs of coincident genes on the same genomic element, there were 17,714 coincident pairs of genes where one is featured on a chromosome and another on a plasmid in the same genome.

### The pangenome as an ecosystem

Almost exclusively, the debate surrounding accessory genome evolution has been framed in terms of the “usefulness” of genes to their host, or the quality of the fit between a particular gene function and the external environment in which the host is found. However, genes also have effects on one another, requiring us to consider higher level conceptualisation of the dynamics of gene gains, losses and the intrinsic and extrinsic forces that shape pangenome evolution. Tansley (1935) developed the theory of the ecosystem, progressing the study of ecology from a focus on individuals to sets of interacting organisms. We here attempt to lay the groundwork to do the same in the context of pangenomes by characterising gene-gene relationships according to their patterns of occurrence. This aids our understanding of gene content evolution by not only showing the extent of intragenomic influences on gene fitness but also how complex relationships can influence gene content evolution both on the scale of the whole accessory genome and for specific examples of sets of genes (McInerney, 2023).

We investigated three signature relationships within the *E. coli* pangenome that we term mutualisms, commensalisms and competition. We have illustrated these relationships using a small subset of the data, outlined in figure 2. By far the most frequent category of relationship is a mutualistic coincident relationship where the joint presence of a pair of genes in genomes is significantly higher than expected from their overall frequency in the dataset. We recovered 20,915 mutualistic co-incident relationships out of our total of 33,138 inferred relationships. Mutualistic relationships are represented in figure 2 (genes A, B, C). Commensal relationships, where one gene, the less abundant of the pair, usually requires the other but the reverse relationship is much weaker or non-existent (see methods) were seen in our networks a total of 2,073 times. Commensal relationships can be seen in figure 2 between gene pairs DA, DB, DC, EA, EB and IH. Finally, competition relationships, which are defined as pairs of genes that avoid each other, were observed in 288 cases.

**Figure 2:**
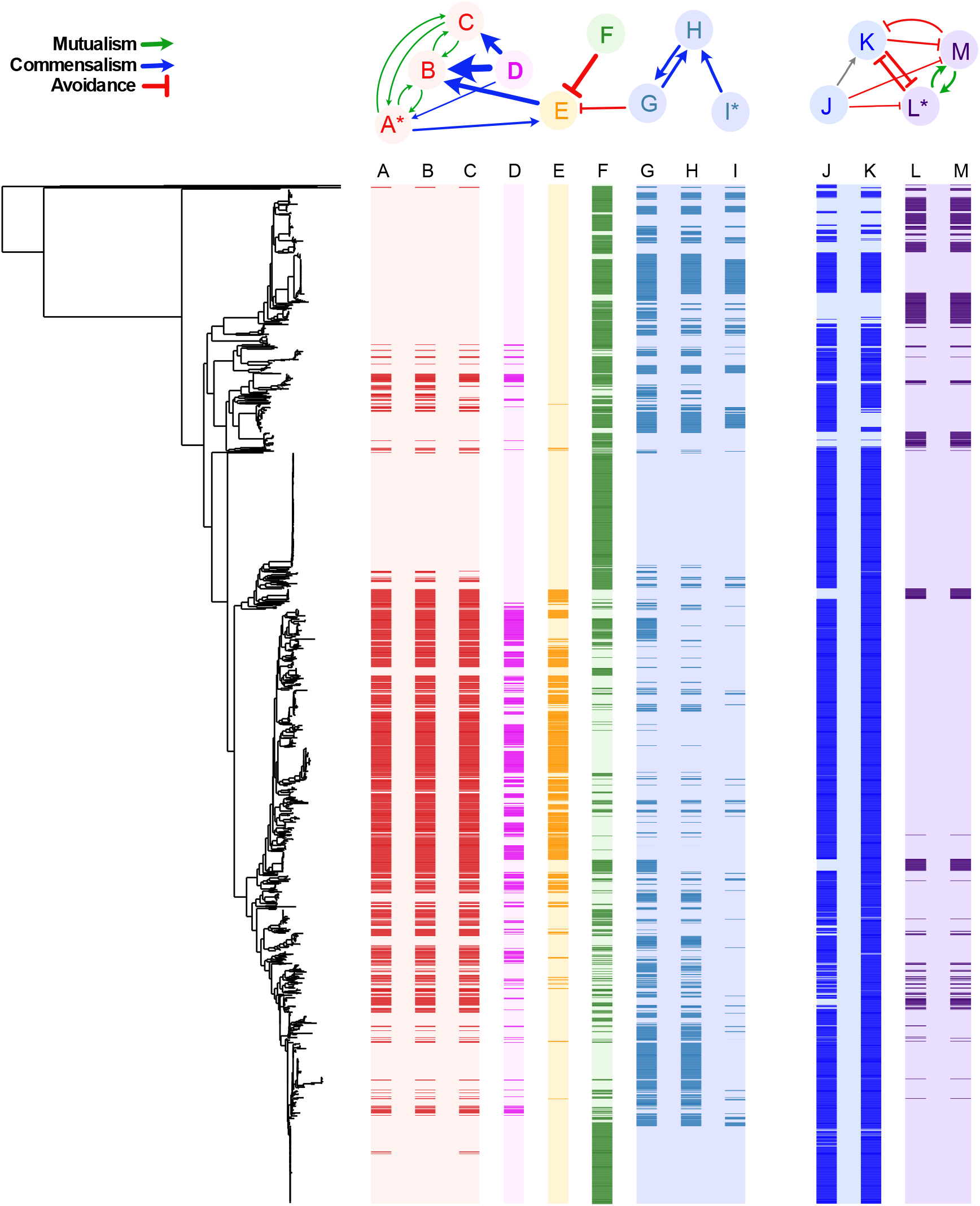
Relationships Between Selected Presence-Absence Patterns in the *E. coli* Pangenome. On top are a network of nodes that represent the presence-absence patterns of the columns directly beneath, as well as the connections between the nodes that represent significant co-occurrence and avoidance relationships. Below left, the backbone phylogeny of the genomes in this study is positioned such that the rows of the heatmap to its right represent the presence or absence of nodes according to the label above each column. In each the colour of presence is indicated by the colour of the text labels in the network and the colour of absence by the background colour of the node. Columns are coloured differently for differentiation rather than their properties. The colours of the arrows in the network indicate the type of association inferred. This figure we produced with aid of a modified version of the program roary_plots.py which is part of the Roary suite of tools (Page et al., 2015). The gene annotations are as follows. Node A: a set of genes with identical presence absence patterns including *farR, hpcG, ttuB, hpcB, hpcE, hpcD, hpcH_2, iolA*, and *hpaB*. Node B: *hpaC*. Node C: *rhaR_2*. Node D: group_39613 (not functionally annotated). Node E: *pac*. Node F: *symE*. Node G: group_13180 (not functionally annotated). Node H: group_19718 (not functionally annotated). Node I: A set of genes with identical presence absence patterns including group_19717, *hsdM*, and *mrr* (see Supplementary Table 4 for full annotations). Node J: *lgoT*. Node K: mdtM. Node L: A set of genes with identical presence absence patterns including group_24769 (*SiaP*), *siaT*, and *nhaK*. Node M: *dctM*:*siaM* (see Supplementary Table 5 for full annotations).

Twenty connected components in our graph consist of genes that show both competition and coincident relationships in which two coincident gene sets have a reduced probability of being in the same genome at the same time (see Figure 2). Although the set of competition relationships is the smallest of our three categories, it represents the interesting situation where one gene makes a genome much less hospitable to another. In figure 2, we see that nodes F and G both predict the absence of node E. The reciprocal is not seen, though an analysis of the importances shows that the reciprocal relationship for E and F is just below our cutoff value (0.00925, when the cutoff was 0.01). To ensure avoidance relationships that we identified are genuine, we carried out a *post hoc* analysis of the avoiding gene pairs and none of the genes share a sequence identity of at least 50% between each family or an E value < 10.

In figure 2 we outline a set of PAPs that represent one or more gene families, that predict the presence or absence of at least one other gene family. In the cases outlined, none are plasmid borne. In addition to being good examples of mutualism, commensalism, and competition, the genes that manifest these PAPs are also of translational importance. For example, PAP E is the “*pac*” gene. During a cell’s response to penicillin, the Pac protein catalyses the hydrolysis of penicillin, forming 6-aminopenicillanate, which is also important in the manufacture of synthetic penicillins (Schumacher et al., 1986, Arroyo et al., 2003). The presence or absence of the encoding gene is predicted accurately in *E. coli* genomes using our Random Forest approach. Using parsimony reconstruction, we estimate that there have been at least 72 changes from present to absent or absent to present for this gene family across the phylogeny. Furthermore, the analysis of its distribution across the tree shows that it has a D score of > 0. Three other PAPs in our dataset are strongly predictive of the presence or absence of *pac*. These are PAPs G (group_13180 (not functionally annotated)) and F (*symE*), which are single gene families, and their presence strongly predicts the absence of *pac*, and the set of genes indicated by node A (*farR, hpcG, ttuB, hpcB, hpcE, hpcD, hpcH_2, iolA*, and *hpaB*) that, conversely, strongly predict the presence of *pac*. Of these three PAPs, perhaps the most striking relationship is its avoidant relationship between PAP F, the gene *symE*, which is more abundant throughout the dataset, but has a genome occupancy pattern that is completely mutually exclusive with *pac* – there are no genomes where both genes are present, although there are several where neither are. *symE* has a parsimony score of 167 and is a translational repressor associated with the SOS response (Kawano et al., 2007).

We also outline the predictive relationships between four other PAPs (Figure 2 J-M). To annotate these genes, a representative protein sequence for each gene family was compared with the NCBI non-redundant protein database using BLASTP. PAP J is annotated as *lgoT*, which is an MFS Transporter. PAP K is annotated as *mdtM*, a multidrug efflux MFS transporter. PAP L is a collection of three distinct families that co-occur perfectly. PAP L includes *nhaK*, a Na+/H+ antiporter, *siaP*, which is a C4-dicarboxylate TRAP transporter substrate-binding protein, and *siaT* which is a TRAP transporter small permease. PAP M is annotated as *dctM:siaM*, which is a TRAP transporter large permease. The presence of PAP J (*lgoT*) is predicted by the presence of PAP K (*mdtM*) (though the reverse is not true). Both PAP J (*lgoT*) and PAP K (*mdtM*) predict the absence of PAPs L and M (collectively, *nhaK, siaP, siaT* and *dctM:siaM*), with all the genes in PAPs L and M predicting each other’s presence.

The most likely observation for any randomly picked genome from this study is that its genome will include either *lgoT* and *mdtM* (both MFS transporters) together (1,882 genomes) or *nhaK, siaP, siaT* and *dctM:siaM* (a sodium ion/proton antiporter and parts of the TRAP transporter complex) together (213 genomes). While this canonical pattern of either one group or the other group, is by far the most likely motif, the data is somewhat noisy, with mdtM co-occurring with at least one gene it normally avoids in 115 genomes. Similarly, the co-occurrence relationships are noisy, for example there are 65 genomes that contain *lgoT* but not *mdtM* compared with the 1,882 that contain both. The most consistent pattern of avoidance is the relationship of *lgoT* with the three genes *siaP, siaT* and *nhaK*, which never co-occur, though there are 23 genomes where none of these genes are present. This means we have found evidence that a multidrug efflux transporter and a Sodium Hydrogen ion antiporter strongly predict the absence of the other. Only 8 genomes contain none of the genes depicted in Figure 2 nodes J-L. All the genes in this set are dynamically lost and gained (parsimony scores: *lgoT*, 71, *mdtM*, 43, *siaP, siaT* and, *nhaK* 57 and *dctM:siaM*, 55), showing that the relationships between these genes have arisen multiple times throughout evolution.

## Discussion

A mechanistic explanation for prokaryotic pangenome origin and evolution is emerging (Domingo-Sananes and McInerney, 2021, McInerney et al., 2020). Here we specifically focussed on detecting pangenome-wide repeated, and predictable patterns of evolution using a Random Forest approach. In effect, we have asked whether gene content evolution is predictable and if within-species evolution is constrained by intragenomic forces.

Though it was necessary to filter the dataset to remove rare and almost universal gene families (almost 50% of the accessory genome), we identified strong predictors for approximately 30% of the remaining gene families. While this leaves much of the accessory genome in a category of non-predictable, given the current data and method of analysis, it must be viewed in the context that we have only used a single species pangenome and therefore, might ask how the inclusion of additional species and datasets might lend additional predictive power. Given that downsampling our dataset reduced the proportion of gene families that could be well predicted, it is likely that the addition of more genomes would aid prediction, and the effect of broadening the taxon sampling would be of interest in future studies. The pattern of repetition and predictability that we observe across more than 2,044 gene families is compatible with a model of deterministic evolution, and more difficult to reconcile with an evolutionary process dominated by contingent events. This compares with those gene families that we were not able to predict accurately for which we cannot rule out contingency on outside factors driving their evolution. Furthermore, given the way in which we have analysed the data, the reasons for observing such widespread predictability stem from intragenomic natural selection causing biases in the cohort of other genes that are acquired, retained, or avoided by a genome. Another possibility is that two (or more) genes are found together because they are both selectively beneficial in the same environment. Hence, when a lineage colonises this environment, both genes are selectively recruited independently. The reasons for co-occurrence and avoidance are largely speculative and would require complex experiments to decipher. Biased presence-absence patterns have been noted previously (Whelan et al., 2020, Mehta et al., 2022, Hall et al., 2021, Whelan et al., 2021).

Throughout this study, we have been using the heuristic that homology is closely related to functional similarity, and that the effects that genes have on one another is due in some way to the encoded functions of the proteins. Furthermore, we assume that these functions remain constant over time, despite evolutionary changes in the gene sequences. This is of course, unlikely to be completely accurate (Koonin, 2005, Stamboulian et al., 2020). Our dataset almost certainly contains genes that are placed into the same family but have different functions and confer different fitness effects. This limitation would certainly weaken our ability to make accurate predictions, though only a small portion of the dataset has verified function.

We note that a lot of gene families are not well predicted given the current dataset and method of analysis. Our dataset might be too small to supply enough power for statistical inference, or indeed genetic background on its own, does not overcome random genetic drift or other evolutionary drivers for some weak fitness effects. Downsampling the dataset produced a significant reduction in prediction power. While we do not know how this will play out with several-fold larger datasets, the direct corelation between dataset size and predictive power suggests that larger datasets should provide predictive capabilities for an increased percentage of the total dataset. Future inclusion of other factors such as external environment, gene expression levels, protein interaction, phenotypic, or modification data, may also aid prediction. Recently, the development of gene-specific evolutionary rate models has shown a significant level of metabolic predictability (Konno and Iwasaki, 2023). Of course, a gene whose presence or absence is non-predictable in the current analysis might continue to be non-predictable in enormous datasets, precisely because it is not impacted by co-occurring genes. Lastly, a complex ensemble of positive and negative intragenomic fitness effects experienced by a gene could result in their non-predictability, even in enormous datasets.

We have taken great care to minimise the confounding effect of genome relatedness. By applying the D-score filter (Fritz and Purvis, 2010), all gene families in our predictive model have a history of being gained or lost at least eight times across the pangenome and furthermore, the distribution of any gene family cannot be “clumped” or restricted to just one part of the backbone tree. This is not a perfect way to eliminate the effect of phylogeny on the associations, but by coupling this approach with a very high threshold for predictability, we end up with an approach that shows corelations that are not just because a pair of genes happen to be in the same clade.

We arbitrarily chose a conservative Gini importance cut-off of 0.01 though there is no standard procedure for choosing a cut-off. Gini importance is a measure of the reduction in ambiguity of the test variable (gene presence or absence), with each node that comprises the prediction variable (a predictor gene), averaged over the trees in the forest. A Gini importance of 0.01 means that the ambiguity of the predicted gene is reduced by an average of 1% in each tree (bearing in mind it will only be sampled in a subset of trees). Gini filtering limited the number of associations to the strongest 33,426 connecting 4,067 nodes, after filtering before filtering by model performance and D-statistic. A weaker cut-off value of 0.005, for example, would have included 70,581 edges connecting 6,830 nodes. The distribution of edge strengths suggests the number of edges will increase exponentially as that threshold is reduced linearly (Supplementary Figure 3). Similarly, we chose an arbitrary, conservative, cut-off of 0.9 for accuracy and F1 score for both classes. Using an 80% accuracy threshold, and F1 score threshold of 0.8 would have included >70% more genes and relationships with the number of predictable genes with D greater than zero almost doubling from 2,044 to 3,704. When we shuffled the genome content data, no genes were predicted to a level close to the thresholds we set, the highest average F score for the two possible states being 0.528, so even some genes that are not classified as predictable deviate from our null expectation.

Linkage clearly plays an important role in coordinating the cooccurrence of sets of genes, but it cannot explain all the results. Also, it is not clear whether linkage is the cause, or a consequence of cooccurrence. According to the selfish operon theory (Lawrence, 1999, Pál and Hurst, 2004) we would expect two genes that provide a fitness benefit when found together in a genome to evolve to be physically closer together on the chromosome, so they are less likely to be separated via recombination events. The most likely explanation is that close linkage is both a cause and effect of being found together. However, in addition to gene family co-occurrence where linkage plays a role, there are thousands of examples of gene families cooccurring in a repeated manner where linkage clearly has not played a role throughout the evolution of the pangenome, either because the genes are separated by a long stretch of DNA or are on different genetic elements.

Genes avoiding being in the same genome is an established phenomenon, with Bruns et al. (2018), for example, showing that within the genus *Salinospora* biosynthetic gene clusters encoding iron siderophores avoid one another. In that case, the clusters encoded iron chelators with very different structures, but near-identical chemical properties, and avoidance most likely stems from functional redundancy. In Figure 2 we outline two situations where avoidance is apparent. In PAPs J-M, both sets of genes encode proton antiporters. It is not clear if the two sets of genes can functionally complement one another, in which case avoidance might be caused by redundancy in the same way as the iron chelators in *Salinosopra*. It is also possible that the functioning of the two antiporters either results in toxic effects when present together or they are competing for the same cellular resources, which would obviously result in a reduction in fitness for the organism that possessed both sets of genes, compared with a close relative that contained only one set of genes. It is also possible that one of the sets of these genes is preferable to the other in some environments, but in others the reverse is true. The situation in Figure 2 PAPs A-I is more enigmatic, with no obvious functional similarity between *pac* and *symE*. However, they are found separately in genomes far more often than expected and furthermore the appearance of one of these genes in a genome occurs simultaneously with the removal of the other and *vice versa* at least 30 times across the phylogeny (Figure 2 PAPs E-F).

Using nomenclature and analogies from ecology, the diversity of motifs embedded in the Random Forest network analysis allows us to identify mutualistic, commensal, and competitive classes of relationship, (see McInerney, 2023). In this sense, the pangenome exists as a broad ecosystem, with individual genomes acting as evolving localities, where genes can either potentiate, or alternatively, reduce the likelihood of the presence of another gene. Just like a macroecological system, we see genes with very high mutualistic co-occurrence patterns, often extending to include several, not just two genes being involved. Focussing on commensal relationships, we identify specific examples of where one or more genes appear to make a genome more hospitable to another or several other genes. In these cases, the more abundant gene is highly likely to arrive in the genome first, or simultaneously with the less abundant gene. We also see many cases where the arrival of a gene into a lineage concomitant with the loss of another gene and this pattern is repeated across the phylogeny.

Ecosystems are known to be dynamic; they also tend to be resilient, and to be somewhat resistant to overall change (Donohue et al., 2016). In our analysis of the *E*. coli pangenome we see features that are consistent with this perspective. We see a very dynamic system of gene gains and losses; we see repetitive gains of the same cohorts of genes, and we see the establishment of sets of relationships that are persistent through time and across the phylogeny. Due to the diversity of the *E. coli* pangenome, each time a gene is recruited to a new genome, it finds itself in a different genetic background, and often the differences are substantial. Despite this variation in genetic background, we observe repeated, predictable patterns of evolution following a gene’s transfer.

With respect to Gould’s “replaying evolution’s tape” thought experiment, our results lead us to suggest that it is likely that rewinding the tape back to the start of *E*. coli evolution would still result in hundreds or thousands of predictable events taking place that are not contingent on highly unlikely events unique to each replaying of the tape. It is unlikely that the exact same evolutionary trajectories would play out, but several motifs would emerge over time.

Other machine learning algorithms such as neural networks may also be able to improve predictions by finding more abstract or subtle patterns in the data that may make a more accurate model. The development of methods, either in the form of additional predictor variables or more sophisticated algorithms, and the application of this method to different bacterial, archaeal, or eukaryotic datasets are enticing future directions which will yield results that help us better understand why pangenomes evolve.

## Supporting information

Supplementary List 1

## Acknowledgements

We would like to thank Mike McDonald for comments and suggestions on this article.

