## Supplementary Figures for "Contingency, Repeatability and Predictability in the Evolution of a Prokaryotic Pangenome"

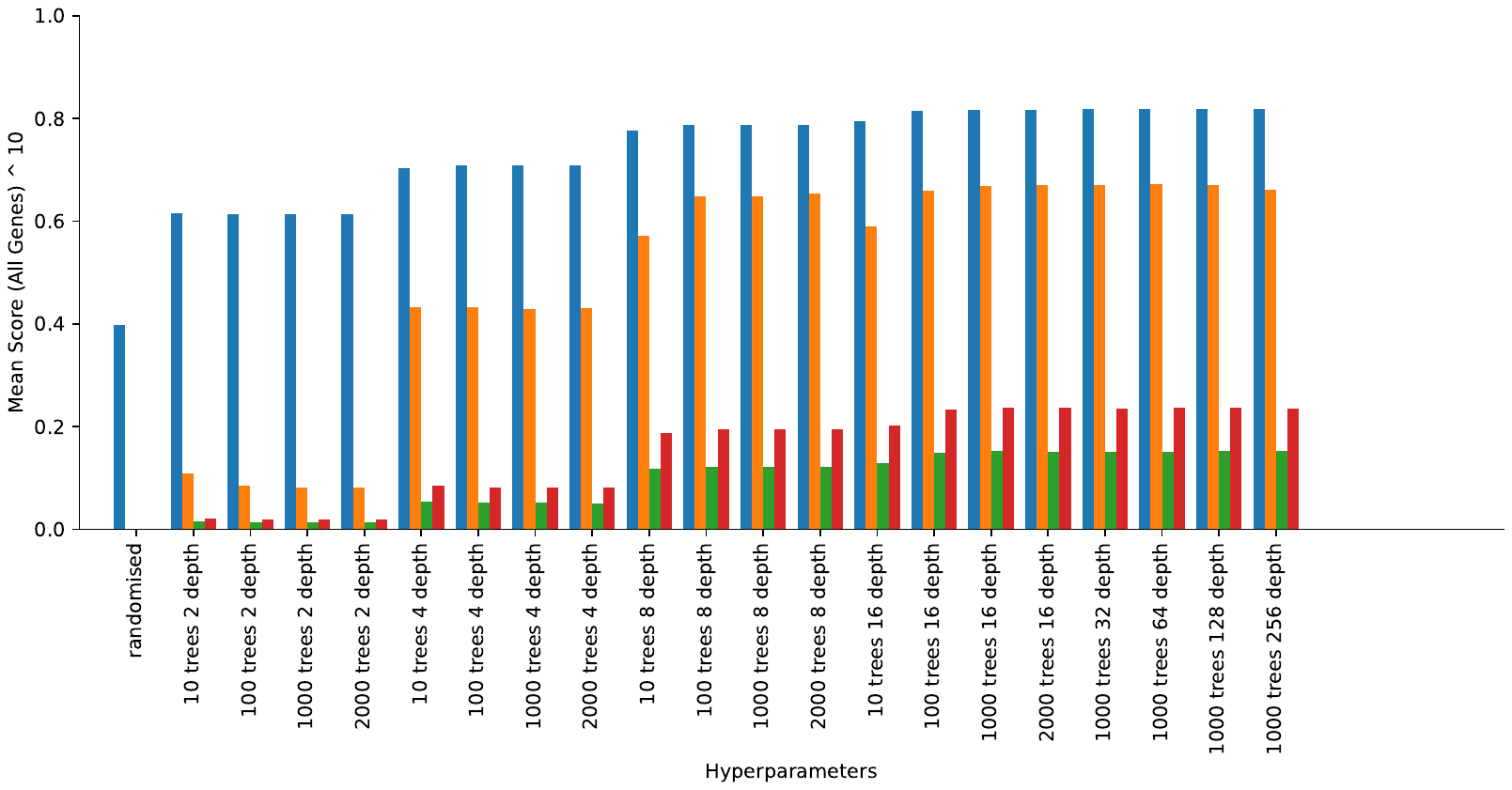

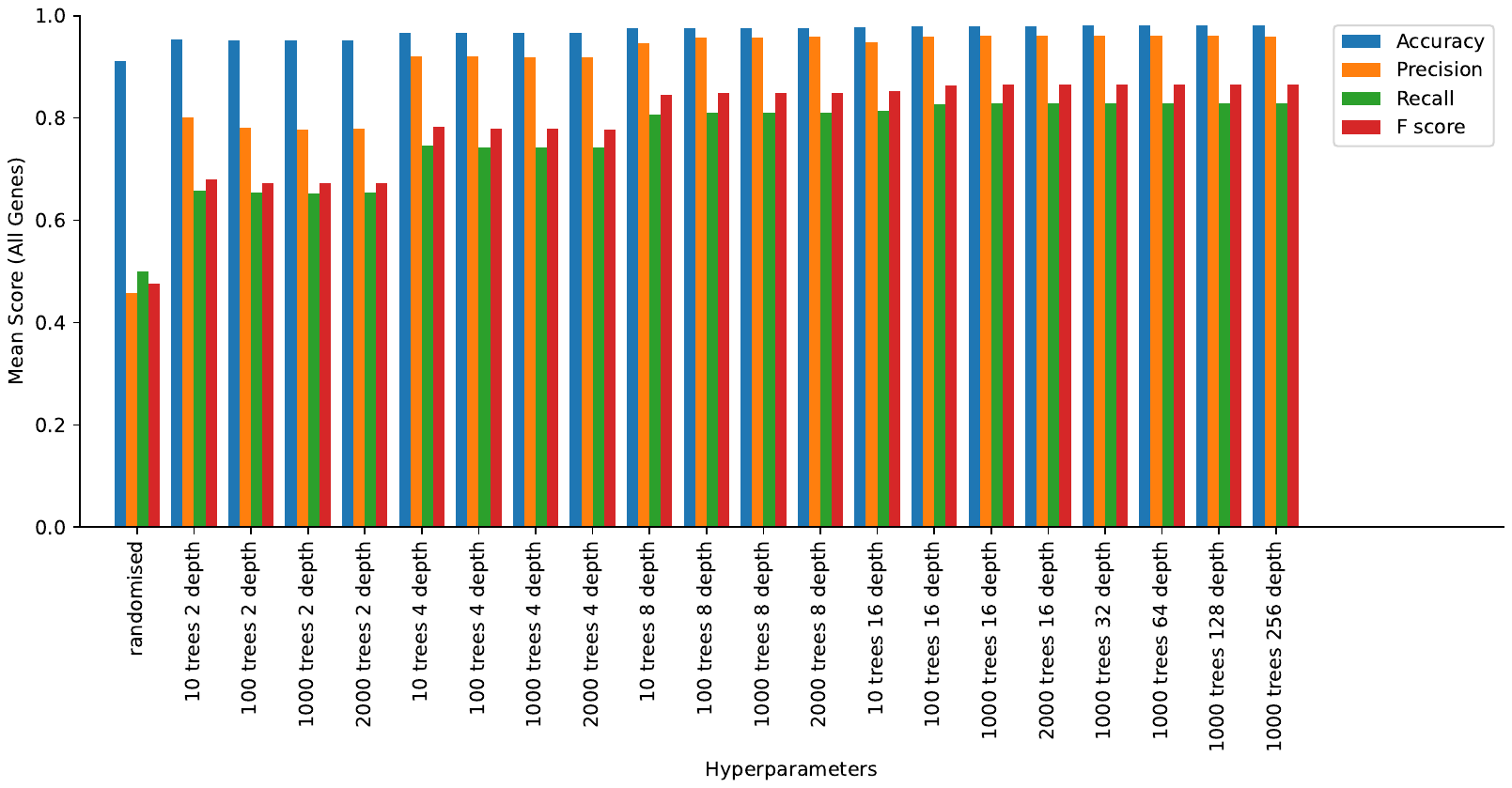

**Supplementary figure 1:** The average Accuracy, Precision, Recall and F-Score for all genes in random forest analyses with varying numbers of trees and maximum tree depth. The values have been transformed to the power of 10 (A) to emphasise the differences between hyperparameter sets.

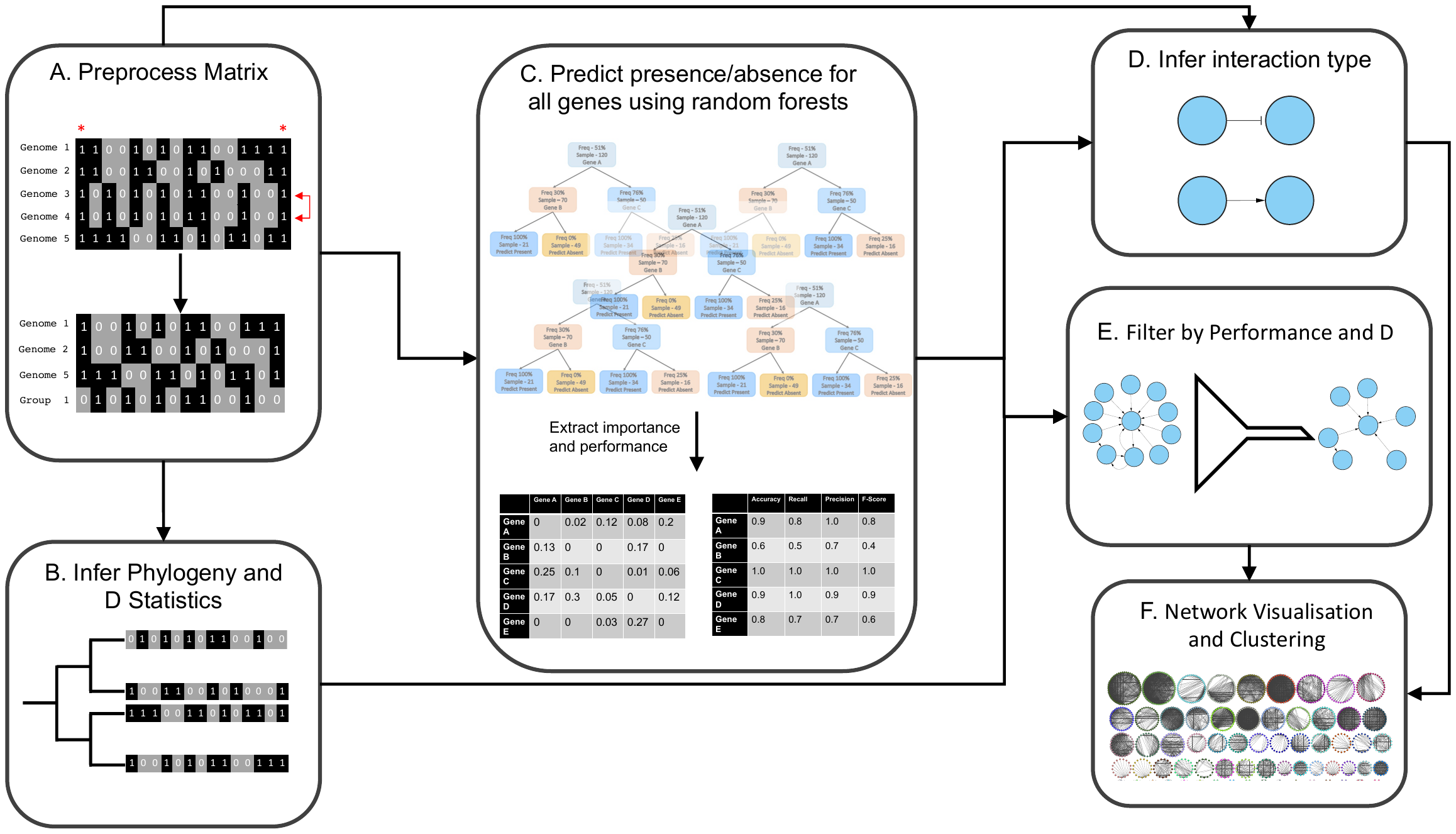

**Supplementary Figure 2:** An illustration of the analytical pipeline taken in this study. The output of each step is used as the input to another step if an arrow points from to it.

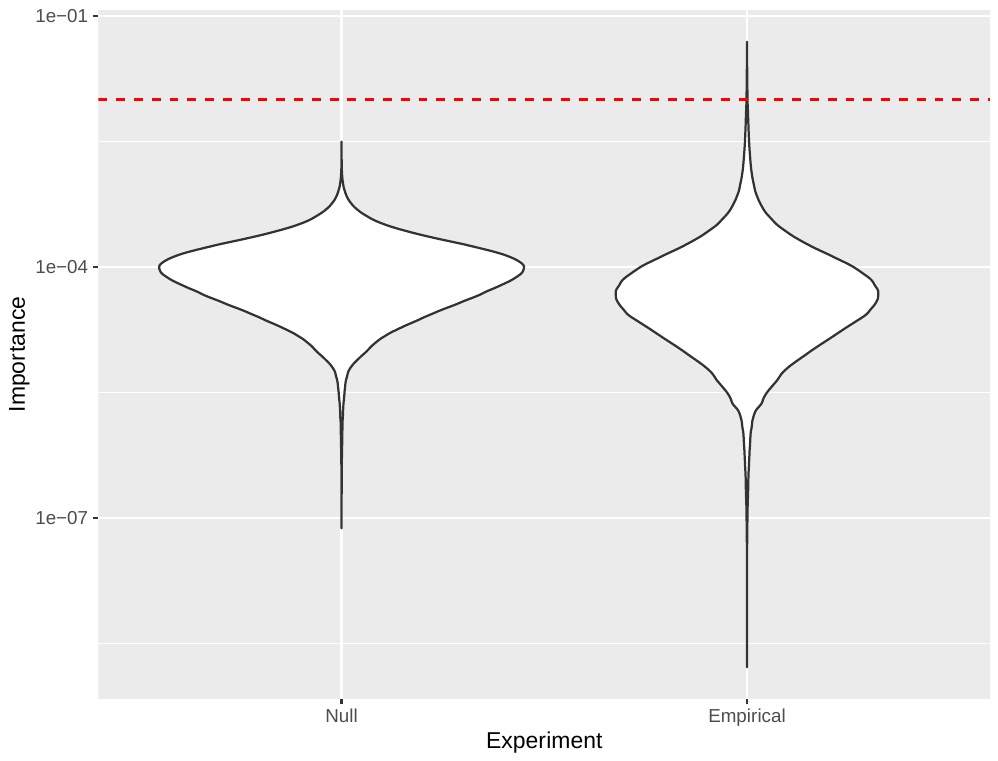

**Supplementary Figure 3**: A comparison the distributions of a random sample (n=1,000,000) of non-0 GINI importance values for each gene in predicting each other gene in the null experiment, where the gene presence absence matrix was randomly shuffled and the empirical analysis where it was not. A red dotted line at y = 0.01 represents the point below which all importance values were discarded. For easier visualisation, the y axis is transformed by a logarithm of base 10.

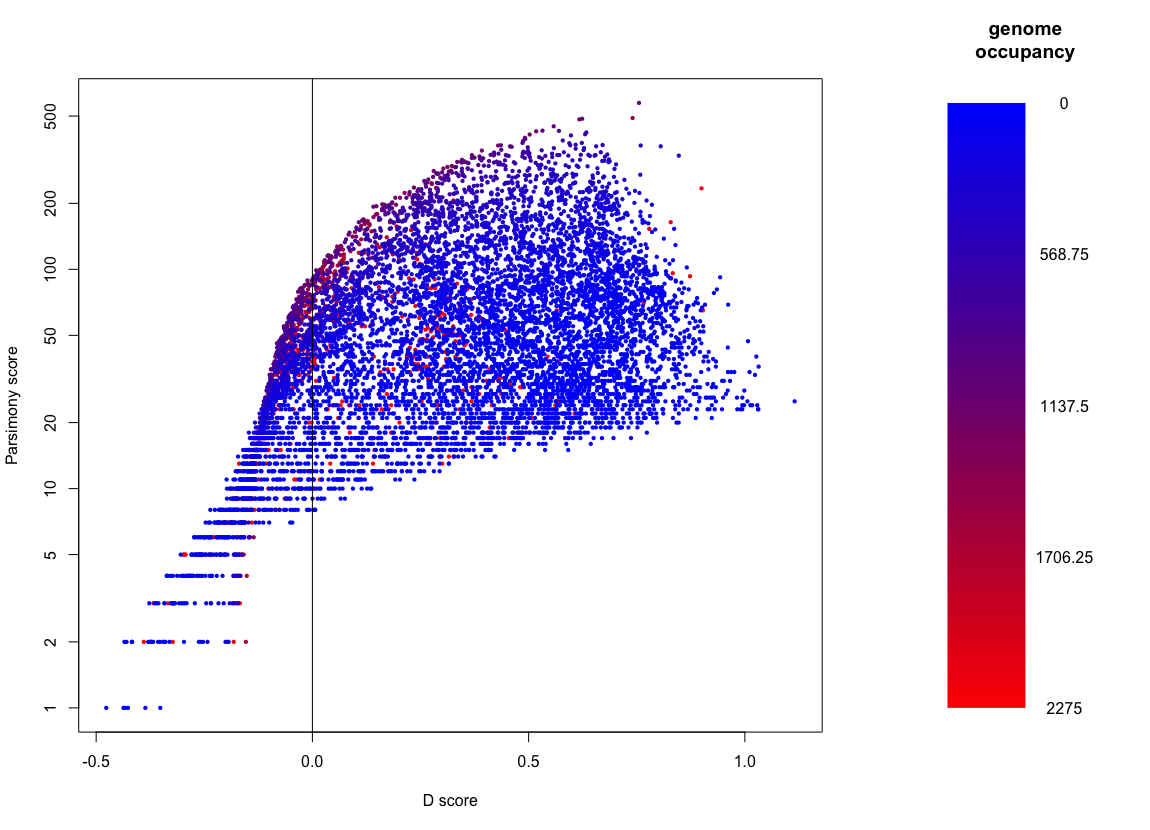

**Supplementary Figure 4:** The relationship between parsimony score (y axis), D score (x axis) and number of genomes in which a gene in present (colour scale). A vertical black line at D score = 0 represents the point at which genes were excluded from our results.

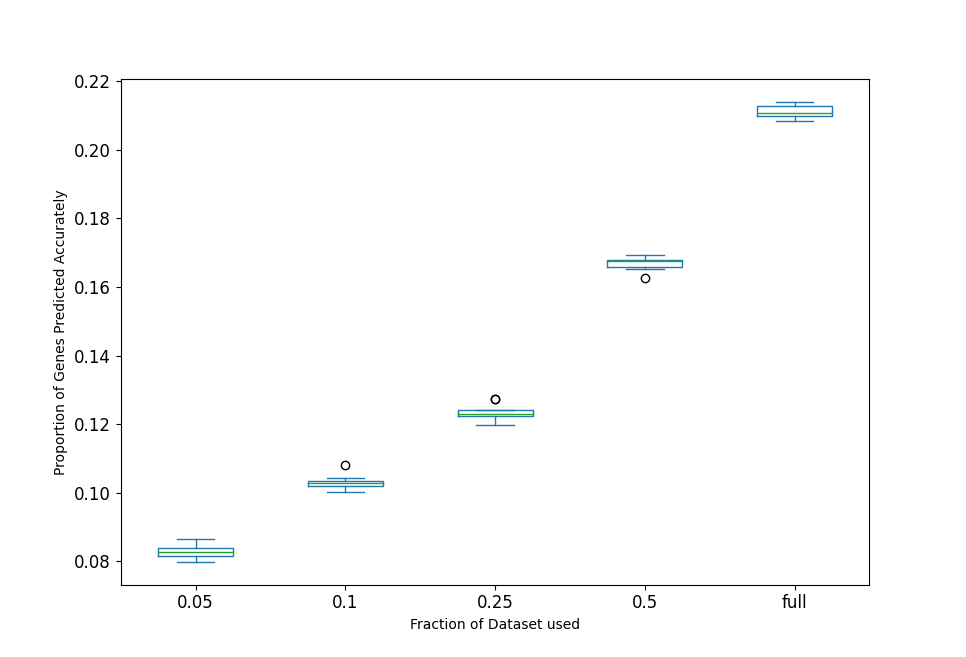

**Supplementary Figure 5:** The proportion of genes predicted accurately after filtering by D-score in a series of experiments where the dataset was down sampled according to the X axis. In each boxplot, the green line represents the median proportion of gene families predicted accurately across 10 repeats, the blue lines between which the box lies are the lower and upper quartiles and the outermost blue horizontal lines are minimum and maximum proportion of genes predicted accurately within1.5 times the interquartile range of the upper or lower quartile. Values outside this range are plotted as circles.

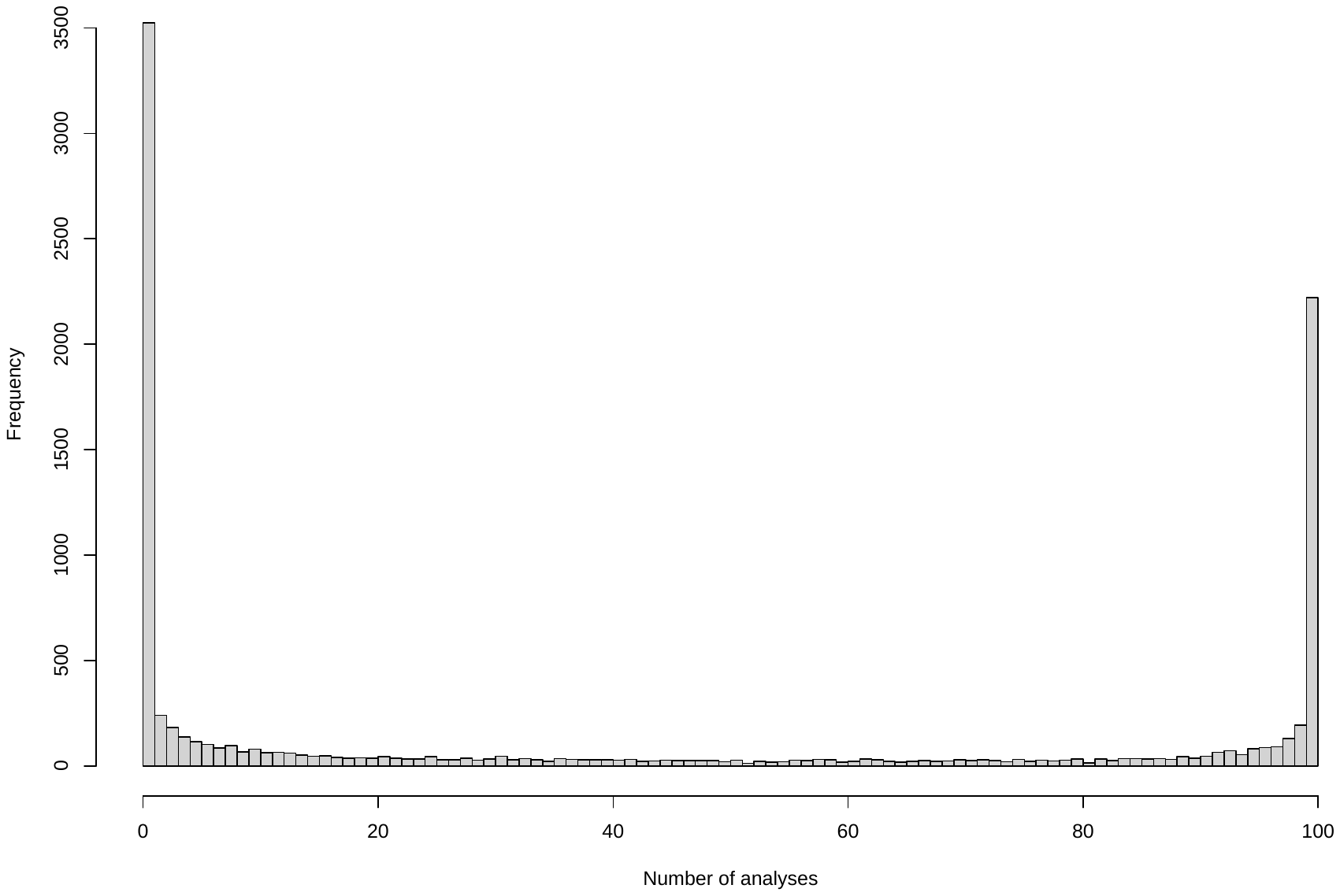

**Supplementary Figure 6:** Histogram showing the number of repeated analyses a given gene is found to be predictable in before filtering for the D score.

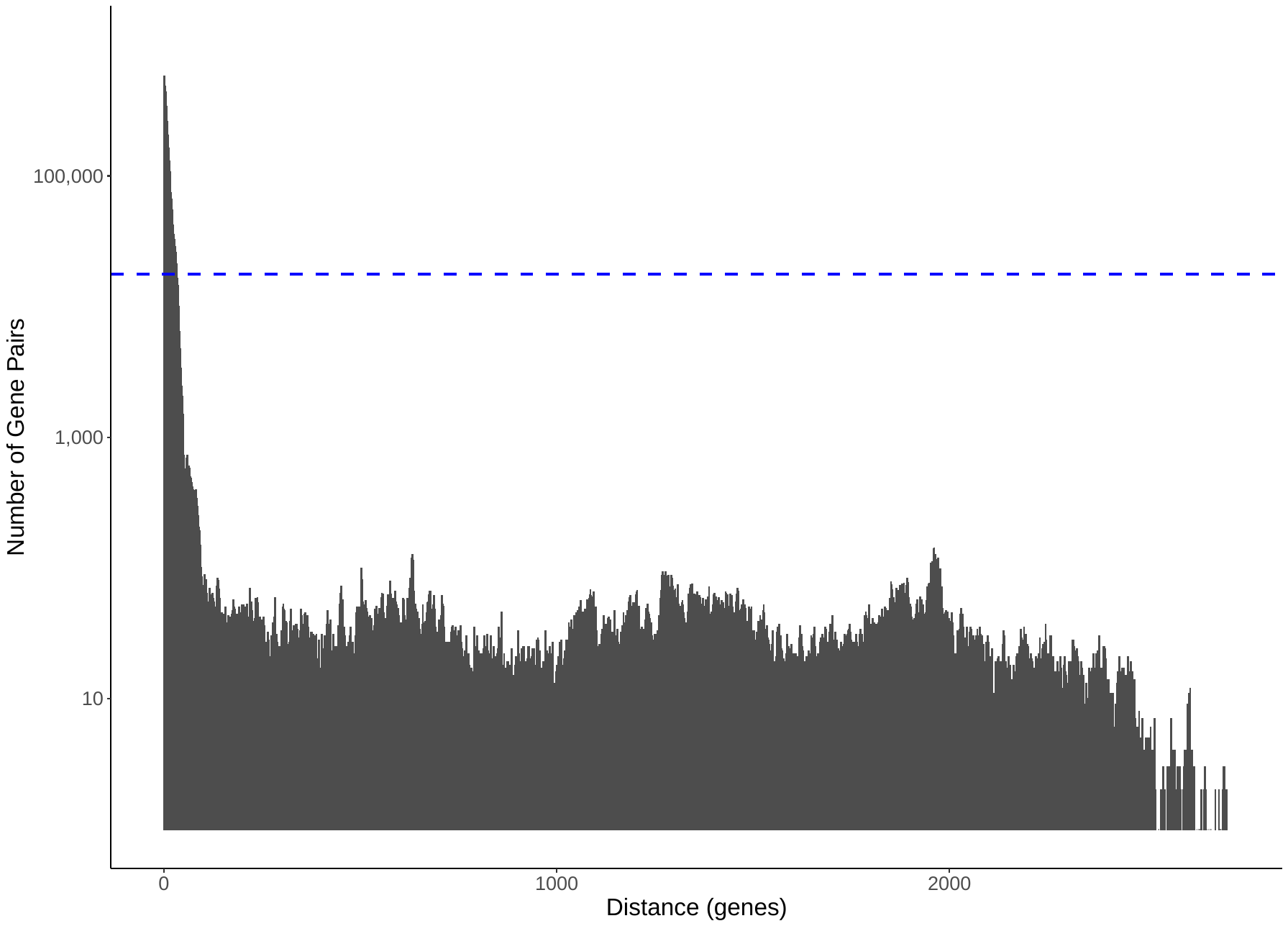
**Supplementary figure 7: Most genes in associated pairs including the most highly predictable genes are closely linked**. The number of associations (log transformed y axis) are plotted against physical distance in number of genes (x axis). A horizontal blue dashed line marks the number of coincident gene pairs in the same genome where each gene occupies a different genomic element (usually chromosome and plasmid).

**Supplementary table 3:** enriched and purified GO terms in the least predictable set of genes.

| **GO** | **NS** | **enrichment** | **name** | **ratio_in_study** | **ratio_in_pop** | **p_uncorrected** | **depth** | **study_count** | **p_fdr_bh** |
| --- | --- | --- | --- | --- | --- | --- | --- | --- | --- |
| **GO:0019634** | BP | e | organic phosphonate metabolic process | 7/870 | 13/12840 | 7.65E-06 | 5 | 7 | 0.000833 |
| **GO:0019700** | BP | e | organic phosphonate catabolic process | 6/870 | 11/12840 | 3.27E-05 | 6 | 6 | 0.00303 |
| **GO:0046434** | BP | e | organophosphate catabolic process | 6/870 | 17/12840 | 0.000615 | 5 | 6 | 0.0383 |
| **GO:0008150** | BP | p | biological_process | 56/870 | 1781/12840 | 1.07E-12 | 0 | 56 | 1.86E-09 |
| **GO:0009987** | BP | p | cellular process | 43/870 | 1420/12840 | 8.42E-11 | 1 | 43 | 7.33E-08 |
| **GO:0044238** | BP | p | primary metabolic process | 16/870 | 733/12840 | 9.36E-09 | 2 | 16 | 5.43E-06 |
| **GO:0006807** | BP | p | nitrogen compound metabolic process | 13/870 | 618/12840 | 1.04E-07 | 2 | 13 | 4.53E-05 |
| **GO:0071704** | BP | p | organic substance metabolic process | 23/870 | 831/12840 | 2.12E-07 | 2 | 23 | 7.39E-05 |
| **GO:0034641** | BP | p | cellular nitrogen compound metabolic process | 10/870 | 513/12840 | 5.71E-07 | 3 | 10 | 0.000136 |
| **GO:0044237** | BP | p | cellular metabolic process | 24/870 | 824/12840 | 6.31E-07 | 2 | 24 | 0.000136 |
| **GO:0043170** | BP | p | macromolecule metabolic process | 13/870 | 581/12840 | 7.06E-07 | 3 | 13 | 0.000136 |
| **GO:0008152** | BP | p | metabolic process | 30/870 | 946/12840 | 7.7E-07 | 1 | 30 | 0.000136 |
| **GO:1901360** | BP | p | organic cyclic compound metabolic process | 10/870 | 505/12840 | 7.82E-07 | 3 | 10 | 0.000136 |
| **GO:0006725** | BP | p | cellular aromatic compound metabolic process | 10/870 | 500/12840 | 1.09E-06 | 3 | 10 | 0.000172 |
| **GO:0046483** | BP | p | heterocycle metabolic process | 10/870 | 492/12840 | 1.5E-06 | 3 | 10 | 0.000217 |
| **GO:0044260** | BP | p | cellular macromolecule metabolic process | 11/870 | 512/12840 | 2.27E-06 | 4 | 11 | 0.000304 |
| **GO:0090304** | BP | p | nucleic acid metabolic process | 8/870 | 431/12840 | 3.32E-06 | 5 | 8 | 0.000412 |
| **GO:0006139** | BP | p | nucleobase-containing compound metabolic process | 10/870 | 470/12840 | 5.48E-06 | 4 | 10 | 0.000636 |
| **GO:0006259** | BP | p | DNA metabolic process | 2/870 | 244/12840 | 1.34E-05 | 6 | 2 | 0.00138 |
| **GO:0044248** | BP | p | cellular catabolic process | 0/870 | 156/12840 | 3.31E-05 | 3 | 0 | 0.00303 |
| **GO:0006310** | BP | p | DNA recombination | 1/870 | 176/12840 | 0.000108 | 7 | 1 | 0.0094 |
| **GO:0061694** | CC | e | alpha-D-ribose 1-methylphosphonate 5-triphosphate synthase complex | 4/870 | 6/12840 | 0.000281 | 5 | 4 | 0.00877 |
| **GO:1904176** | CC | e | carbon phosphorus lyase complex | 4/870 | 8/12840 | 0.00118 | 3 | 4 | 0.026 |
| **GO:0061695** | CC | e | transferase complex, transferring phosphorus-containing groups | 4/870 | 10/12840 | 0.00316 | 4 | 4 | 0.0493 |
| **GO:0110165** | CC | p | cellular anatomical entity | 39/870 | 1254/12840 | 3.59E-09 | 1 | 39 | 5.61E-07 |
| **GO:0005575** | CC | p | cellular_component | 49/870 | 1371/12840 | 1.04E-07 | 0 | 49 | 8.11E-06 |
| **GO:0016020** | CC | p | membrane | 18/870 | 634/12840 | 1.03E-05 | 2 | 18 | 0.000537 |
| **GO:0071944** | CC | p | cell periphery | 18/870 | 616/12840 | 2.41E-05 | 2 | 18 | 0.00094 |
| **GO:0005622** | CC | p | intracellular anatomical structure | 14/870 | 451/12840 | 0.000779 | 2 | 14 | 0.0203 |
| **GO:0005737** | CC | p | cytoplasm | 14/870 | 440/12840 | 0.00133 | 2 | 14 | 0.026 |
| **GO:0005886** | CC | p | plasma membrane | 15/870 | 456/12840 | 0.00162 | 3 | 15 | 0.028 |
| **GO:0003674** | MF | p | molecular_function | 41/870 | 1253/12840 | 2.12E-08 | 0 | 41 | 2.11E-05 |
| **GO:0003824** | MF | p | catalytic activity | 20/870 | 771/12840 | 1.7E-07 | 1 | 20 | 8.45E-05 |
| **GO:0140640** | MF | p | catalytic activity, acting on a nucleic acid | 1/870 | 177/12840 | 0.00011 | 2 | 1 | 0.0364 |

**Supplementary table 4:** The names of genes inferred by Panaroo, their labels in the main text and the node they correspond to (figure 2 main text)

| Gene (group) | Name in main text | Full Panaroo annotation |
| --- | --- | --- |
| A | *farR* | farR |
| A | *hpcG* | hpcG |
| A | *ttuB* | ttuB_1~~~ttuB_2~~~ttuB |
| A | *hpcB* | hpcB~~~hpcB_1~~~hpcB_2 |
| A | *hpcE* | hpcE~~~hpcE_1~~~hpcE_2 |
| A | *hpcD* | hpcD |
| A | *hpcH* | hpcH_2~~~hpcH~~~hpcH_1 |
| A | *iolA* | iolA~~~xylG_2~~~betB_1~~~xylG_1~~~tgnC~~~aldB_1~~~aldB_2~~~betB_2 |
| A | *hpaB* | hpaB_2~~~hpaB~~~hpaB_1 |
| B | *hpaC* | hpaC |
| C | *rhaR* | rhaR_2~~~rhaR_4~~~rhaR_3~~~rhaR_5 |
| D | group_39613 | group_39613 |
| E | *pac* | pac_1~~~pac_3~~~pac~~~pac_2 |
| F | *symE* | symE~~~symE_3~~~symE_8~~~symE_1~~~symE_2~~~symE_4 |
| G | group_13180 | group_13180 |
| H | group_19718 | group_19718 |
| I | *hsdM* | hsdM |
| I | *mrr* | Mrr |

Supplementary table 6: The names of genes inferred by Panaroo and how we refer to them in the main text

| Gene (group) | Name in text | Name given by Panaroo |
| --- | --- | --- |
| J | *lgoT* | lgoT_1~~~lgoT~~~lgoT_2 |
| K | *mdtM* | mdtM~~~mdtM_2~~~mdtM_1~~~mdtL_1 |
| L | *nhaK* | nhaK_2~~~nhaK_1 |
| L | *siaP* | group_24769 |
| L | *siaT* | siaT~~~siaT_2 |
| M | *dctM:siaM* | dctM~~~dctM_2~~~dctM_1~~~siaM_2~~~siaM_1 |
